# The ERAD ubiquitin ligase Doa10 negatively regulates Atg32-mediated mitophagy

**DOI:** 10.1101/2024.04.08.588568

**Authors:** Akinori Eiyama, Mashun Onishi, Hana Štupica, Yukiko Omi, Sachiyo Nagumo, Kunio Nakatsukasa, Koji Okamoto

## Abstract

Degradation of defective or superfluous mitochondria via mitophagy, a specialized form of selective autophagy, is important for maintaining mitochondrial quality and quantity. In yeast, the pro-mitophagic factor Atg32 is transcriptionally upregulated upon oxidative stress and anchored to the mitochondrial surface, where it acts as a molecular signal to initiate efficient degradation of mitochondria. However, how the protein levels of Atg32 are regulated post-translationally remains enigmatic. Here we show that the endoplasmic reticulum (ER) serves as a hub to govern Atg32 protein turnover. We found that the ER-associated degradation (ERAD) E3 ligase Doa10 interacts with Atg32, leading to its degradation by the proteasome. Furthermore, we show that Atg32 is destined for the ER in a manner dependent on the GET (guided entry of tail-anchored proteins) pathway, which mediates the delivery of tail-anchored (TA) proteins to the ER where Atg32 is potentially recognized by Doa10. Notably, Doa10 deficiency increased Atg32 levels and enhanced mitophagy under respiratory conditions, thus determining that ERAD serves as a brake on mitophagy.

## INTRODUCTION

Mitochondria-specific autophagy, termed mitophagy, is one of the catabolic pathways conserved from yeast to humans, whose failure can be linked to several pathological consequences (Palikaras et al, 2018; Onishi et al, 2021; Uoselis et al, 2023). In this process, mitochondria are sequestered into double-membrane vesicles called autophagosomes and transported to the lysosome (in mammals) or the vacuole (in yeast) for degradation. In the budding yeast *Saccharomyces cerevisiae*, Atg32, a single-pass transmembrane protein on the outer mitochondrial membrane (OMM), acts as an essential mitophagy receptor and “eat me” signal to initiate sequestration of mitochondria by autophagosomes (Okamoto et al, 2009; Kanki et al, 2009). Atg32 is induced at the transcriptional level in response to oxidative stress (Okamoto et al, 2009; Eiyama et al, 2015). In addition to transcriptional upregulation, post-translational modification of Atg32 is required to modulate Atg32 activity and mitophagy. Atg32 is phosphorylated in a manner dependent on casein kinase 2 under mitophagy-inducing conditions, which increases the affinity of Atg32 for Atg11, a scaffold protein that recruits other Atg proteins to the mitochondrial surface (Kanki et al, 2009, 2013; Okamoto et al, 2009; Aoki et al, 2011; Kondo-Okamoto et al, 2012). Conversely, Atg32 dephosphorylation is mediated by the PP2A-like phosphatase Ppg1. Ppg1 is dually localized on the ER and mitochondria, and suppresses Atg32 phosphorylation, Atg32–Atg11 interactions, and mitophagy (Furukawa et al, 2018; Innokentev et al, 2020; Onishi et al, 2023). Loss of Atg32 almost completely abolishes mitophagy, whereas its overexpression enhances mitophagic degradation, suggesting that Atg32 protein levels are one of the rate-limiting factors that determine the number of mitochondria to be degraded (Okamoto et al, 2009; Kanki et al, 2009; Eiyama et al, 2015).

Although accumulating evidence suggests that defects in mitophagy are associated with pathophysiological conditions such as neurodegenerative and cardiovascular diseases, overactivated basal mitophagy in mammals could also be detrimental (Nguyen-Dien et al, 2023; Cao et al, 2023; Elcocks et al, 2023; Chen et al, 2023; Niemi et al, 2023; Sun et al, 2024). For example, loss of the SCF-FBXL4 ubiquitin E3 ligase complex, which ubiquitinates and targets mammalian mitophagy receptors for degradation, accelerates basal mitophagy. Deficiency of this E3 ligase complex appears to be associated with reduced mitochondrial content such as mitochondrial DNA and mitochondrial respiratory defects (Nguyen-Dien et al, 2023; Cao et al, 2023; Elcocks et al, 2023). These findings suggest that in addition to mechanisms that initiate mitophagy, cells also need to have a brake control system that limits mitophagic activity to finely regulate cellular homeostasis.

ERAD plays a central role in the quality control mechanisms of the ER, where non-native and misfolded proteins are ubiquitinated and degraded to prevent their abnormal accumulation and aggregation in the ER (Vembar et al, 2008; Ruggiano et al, 2014; Nakatsukasa et al, 2015; Wu et al, 2018; Bhattacharya et al, 2019; Christianson et al, 2022). ERAD involves multiple steps; substrate recognition by molecular chaperones and accessory proteins, retrotranslocation of target substrates to the cytosol, ubiquitination of substrates and subsequent degradation by the proteasome. In the process of ERAD in yeast, two major ERAD E3 ligases mediate the ubiquitination of target substrates: Doa10, which targets proteins with misfolded domains in the cytoplasmic side of the ER membrane (ERAD-C), and Hrd1, which targets proteins with misfolded domains in the ER lumen (ERAD-L) or intramembrane (ERAD-M) (Bordallo et al, 1998; Hill et al, 2000; Bays et al, 2001; Vashist and Ng, 2004; Carvalho et al, 2006; Nakatsukasa et al, 2008). Originally well-characterized as a pathway for degrading misfolded proteins to ensure protein homeostasis in the ER, accumulating studies also suggest that many branches of ERAD appear to operate protein turnover of a wider range of substrates and are involved in ER- and non-ER-related functions (Ruggiano et al, 2016; Zhou et al, 2020; Ji et al, 2023). In particular, ERAD deficiency is associated with mitochondrial dysfunction in several tissues and cells, further highlighting the potential role of ERAD in mitochondrial integrity (Hu et al, 2019; Liu et al, 2020).

In this study, we show that ERAD acts to reduce Atg32 protein levels. We also found that a fraction of Atg32 is localized to the ER via the GET pathway, which mediates the insertion of TA proteins into the ER membrane, and interacts with Doa10. We further demonstrate that Atg32 is ubiquitinated, and that its protein stability is increased by inhibition of the proteasome. Remarkably, loss of Doa10 increases Atg32 and promotes mitophagy under respiratory conditions. Taken together, our data suggest that Doa10-mediated downregulation of Atg32 protein levels contributes to limiting mitochondrial degradation.

## RESULTS

### The Doa10 ERAD E3 ligase restricts Atg32 protein levels and mitophagy

To investigate whether ERAD is involved in Atg32-mediated mitophagy, we first examined whether Atg32 protein levels are altered in cells lacking ERAD components. Cells lacking Doa10 showed increased Atg32 protein levels, whereas loss of Hrd1 did not significantly alter the expression profile of Atg32 (Fig. 1A). Doa10 forms a complex with the E2 enzymes Ubc6 and Ubc7 (Chen et al, 1993; Ruggiano et al, 2014; Weber et al, 2016). Consistently, Ubc6 or Ubc7-deficient cells also exhibited higher levels of Atg32 protein than wild-type cells (Fig. 1B). These data suggest that Doa10-mediated ERAD acts to reduce Atg32 protein levels.

**Figure 1.**
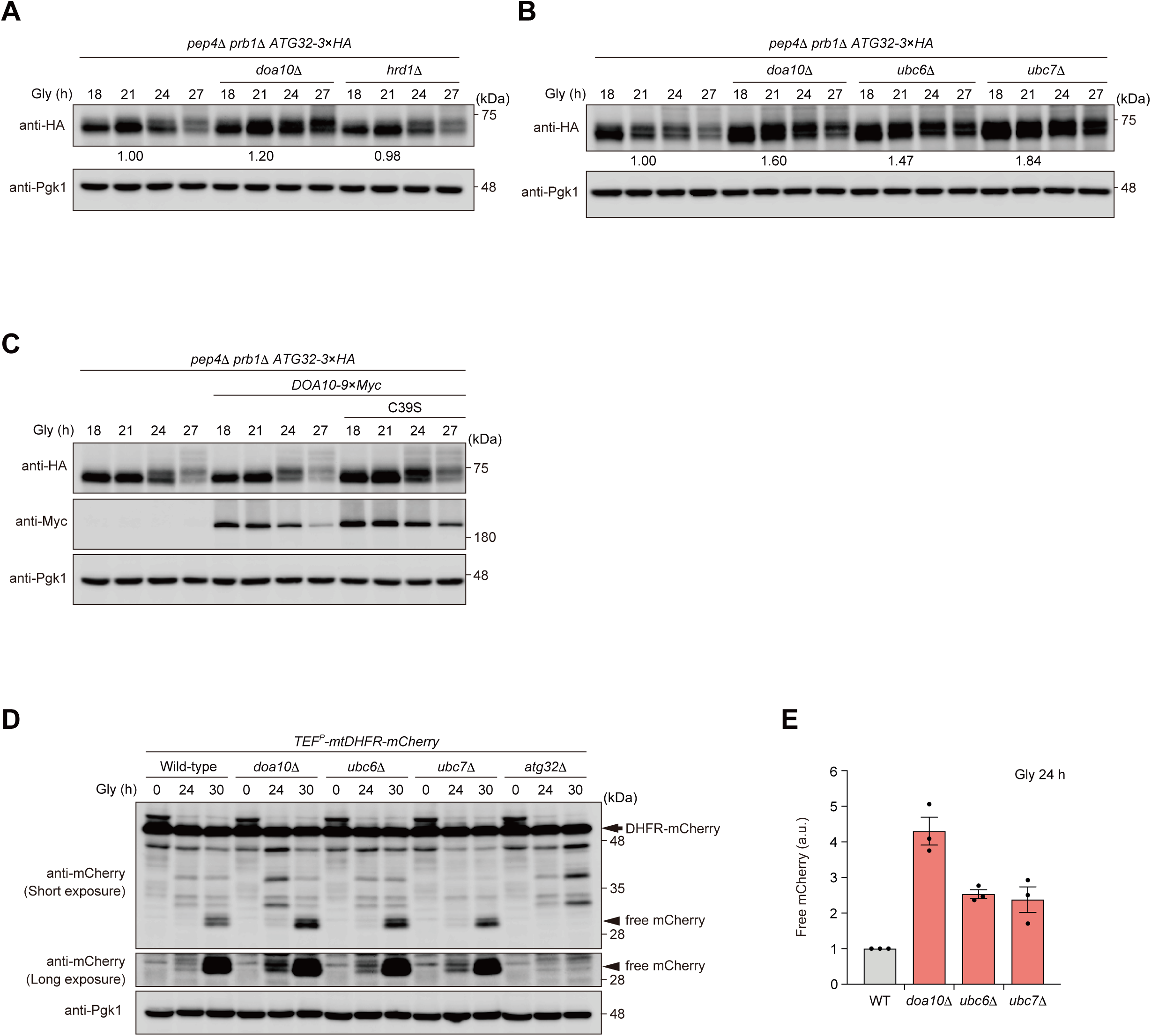
The Doa10 ERAD E3 ligase restricts Atg32 protein levels and mitophagy. **(A)** Atg32-3×HA-expressing wild-type, *doa10*Δ and *hrd1*Δ cells were grown in glycerol medium (Gly), collected at the indicated time points, and subjected to western blotting. Numbers indicate the peak intensities of signals of Atg32-3×HA. **(B)** Atg32-3×HA-expressing wild-type, *doa10*Δ*, ubc6*Δ and *ubc7*Δ cells were grown in glycerol medium (Gly), collected at the indicated time points, and subjected to western blotting. Numbers indicate the peak intensities of signals of Atg32-3×HA. **(C)** The endogenous Atg32-3×HA-expressing cells harboring Doa10 wild-type or a Doa10^C39S^ mutant were grown in glycerol medium (Gly), collected at the indicated time points, and subjected to western blotting. **(A-C)** All strains are *pep4 prb1*-double-null derivatives defective in vacuolar degradation of Atg32-3×HA via mitophagy. Pgk1 was monitored as a loading control. **(D)** Mitochondria-targeted DHFR-mCherry-expressing (mito-DHFR-mCherry) wild-type, *doa10*Δ, *ubc6*Δ, *ubc7*Δ, and *atg32*Δ cells were grown for the indicated time points in glycerol medium (Gly), and subjected to western blotting. Generation of free mCherry indicates transport of the maker to the vacuole. Pgk1 was monitored as a loading control. **(E)** The amounts of free mCherry in cells analyzed in **(D)** were quantified in three experiments. The signal intensity value of free mCherry in wild-type cells at the 72 h time point was set to 100%. Data represent the averages of all experiments (*n* = 3 independent cultures, means ± s.e.m.).

To examine whether the ubiquitin ligase activity of Doa10 modulates Atg32 protein levels, we generated a Doa10 variant lacking its enzymatic activity. A previous study reported that a serine substitution at Cys-39 (C39S) of Doa10 leads to loss of its enzymatic activity (Swanson et al, 2001). We therefore monitored Atg32 protein levels in cells expressing Myc-tagged Doa10^C39S^. We found that Atg32 is increased in cells expressing a Doa10^C39S^-3×Myc variant (Fig. 1C). Thus, we concluded that Atg32 protein levels are reduced in a manner dependent on the ubiquitin ligase activity of Doa10.

We next asked whether loss of Doa10 may likewise promote mitophagy by accumulating Atg32. To test this possibility, we performed mitophagy assays using the mitochondrial matrix-localized DHFR (dihydrofolate reductase)-mCherry (mito-DHFR-mCherry) probe (Eiyama et al 2017, Nagumo et al 2017; Calvelli et al, 2020). When mitochondria are transported to the vacuole, mito-DHFR-mCherry is processed by vacuolar proteases to generate free mCherry, allowing semi-quantitative detection of mitochondrial degradation. Mitophagy in yeast is induced when cells are grown in media containing non-fermentable carbon sources such as glycerol, and start to accumulate respiratory active mitochondria. In those cells that reach the post-log phase under respiratory conditions, a substantial fraction of mitochondria are transported to the vacuole and degraded via mitophagy. We found that mitophagy was enhanced in cells lacking Doa10, Ubc6 or Ubc7 in the early phase of respiratory growth (Fig. 1D and E). In addition, overexpression of Atg32 enhanced mitophagy under respiratory conditions (Eiyama et al, 2015) (Fig. S1A and B), together supporting the idea that increased Atg32 protein levels in *doa10*-null cells can lead to enhanced mitophagy.

We further sought to clarify whether loss of Doa10 affects the other autophagy-related events. First, we monitored the processing of Tdh3 (triosephosphate dehydrogenase 3, a major GAPDH isoenzyme in yeast)-mCherry, a fluorescent marker highly expressed in the cytosol. Upon autophagy induction, Tdh3-mCherry is non-selectively sequestered into autophagosomes, transported to the vacuole, and processed to generate free mCherry dependently on Atg7, a protein essential for autophagy (Mizushima et al, 1998). The accumulation of free mCherry was not significantly altered in cells lacking Doa10 under respiratory conditions (Fig. S2A and B), suggesting that Doa10 is dispensable for bulk autophagy.

Second, we examined the cytoplasm-to-vacuole targeting (Cvt) pathway, a selective autophagy-related process where vacuolar enzymes such as the amino peptidase Ape1 are transported from the cytosol to the vacuole (Shintani et al, 2002; Suzuki et al, 2002). Atg19 mediates the transport of precursor Ape1 (prApe1) to the vacuole and leads to Ape1 processing to generate mature Ape1 (mApe1) (Scott et al, 2001). This processing of Ape1 was detectable in *doa10*-null cells under respiratory conditions (Fig. S2C), indicating that loss of Doa10 slightly or hardly affects the Cvt pathway.

Third, we examined peroxisome-specific autophagy (pexophagy) by monitoring Pot1-mCherry, a peroxisomal matrix-localized marker transported to the vacuole in a manner dependent on Atg36 (Motley et al, 2012) and generates free mCherry. We found that *doa10*-null cells underwent degradation of peroxisomes with higher efficiency (135% compared to wild-type cells at the 72 h time point) (Fig. S2D and E), raising the possibility that Doa10 may act, at least to some extent, in pexophagy.

Similarly, ER-specific autophagy (ER-phagy) under respiratory conditions was analyzed by monitoring Sec63-mCherry, a marker localized in the ER, and transported to the vacuole and processed to generate free mCherry in an Atg39- and Atg40-dependent manner, two proteins essential for ER-phagy (Mochida et al, 2015) Under the same conditions, the processing of Sec63-mCherry was not significantly altered by loss of Doa10 (Fig. S2F and G), supporting the idea that the Doa10 complex may play a minor role, if any, in ER-phagy.

### Atg32 is ubiquitinated under respiratory conditions

Although we observed that the E3 ubiquitin ligase activity of Doa10 modulates Atg32 protein levels (Fig. 1A, B and C), it remains controversial whether Atg32 is degraded by the ubiquitin-proteasome system (UPS). A previous study has reported that Atg32 stability is regulated independently of the proteasome (Levchenko et al, 2016), while a study from another group has observed proteasome-mediated degradation of Atg32 (Camougrand et al, 2020). We therefore sought to define whether the UPS is involved in Atg32 protein turnover under mitophagy-inducing non-fermentable conditions. To address this point, cells are treated with the proteasome inhibitor MG132. These cells are depleted with Pdr5, a multidrug-exporting ATP-binding cassette transporter (Prasad et al, 2012), maximizing the accumulation and efficiency of MG132 to inhibit the UPS in these cells. Treatment with MG132 suppressed the reduction of Atg32 compared to cells treated with DMSO (Fig. 2A), indicating that Atg32 turnover is partially mediated by the proteasome. We further confirmed that Atg32 is slightly stabilized upon Doa10 loss in cycloheximide (CHX)-treated cells (Fig. 2B).

**Figure 2.**
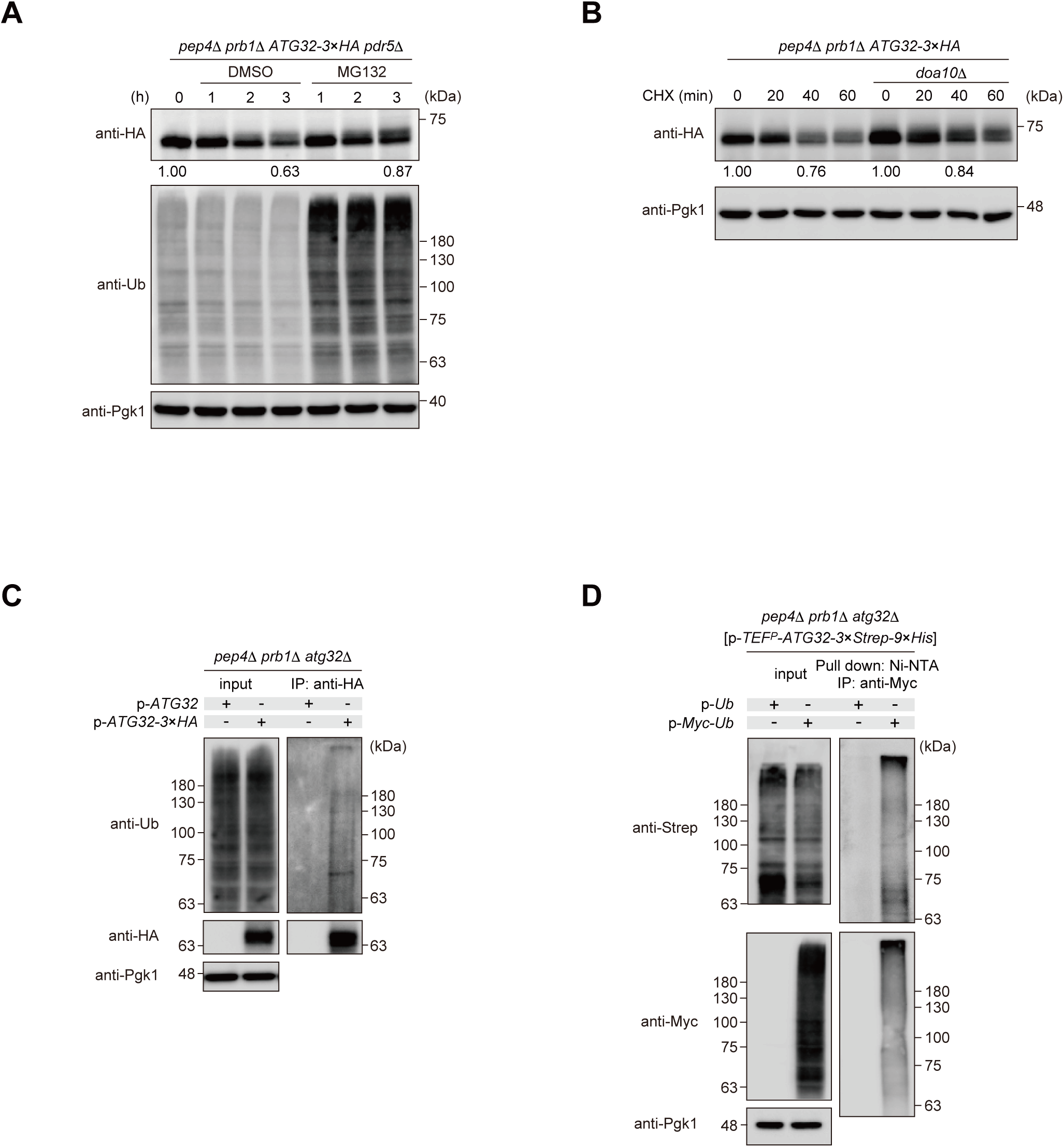
Atg32 is ubiquitinated under respiratory conditions. **(A)** The endogenous Atg32-3×HA-expressing cells lacking Pdr5 were treated with DMSO or MG132, collected at the indicated timepoints, and subjected to western blotting. Pgk1 was monitored as a loading control. Numbers indicate the peak intensities of signals of Atg32-3×HA. **(B)** The endogenous Atg32-3×HA-expressing wild-type cells and *doa10*Δ cells were grown in glycerol medium (Gly), treated with cycloheximide (CHX), collected at the indicated time points, and subjected to western blotting. Pgk1 was monitored as a loading control. Numbers indicate the peak intensities of signals of Atg32-3×HA. **(C)** *atg32*Δ cells expressing chromosome- or plasmid-encoded versions of Atg32 or Atg32-3×HA were grown in glycerol medium, and subjected to coimmunoprecipitation using anti-HA antibody-conjugated agarose. Eluted immunoprecipitates (IP) and detergent-solubilized mitochondria-enriched fractions (input) were analyzed by western blotting. Pgk1 was monitored as a loading control. **(D)** Myc-tagged or non-tagged ubiquitin-expressing *atg32*Δ cells were transfected with a plasmid encoding ATG32-3×Strep-9×His under a TEF promoter. These cells were grown in glycerol medium and subjected to pull-down assay using Ni-NTA, and subjected to coimmunoprecipitation using anti-Myc antibody-conjugated agarose. Pgk1 was monitored as a loading control. **(A-D)** All strains are *pep4 prb1*-double-null derivatives defective in vacuolar degradation of Atg32-3×HA via mitophagy.

These findings prompted us to determine whether Atg32 is ubiquitinated under respiratory conditions. To this end, we performed coimmunoprecipitation (Co-IP) assays using cells expressing HA-tagged Atg32. We observed that ubiquitin was coprecipitated with Atg32-3×HA (Fig. 2C), indicating that ubiquitin interacts with Atg32. To exclude the possibility that these results reflect the ubiquitination of Atg32-interacting partners rather than Atg32 itself, we performed a tandem affinity purification (TAP) using cells expressing Strep and His-tagged Atg32 and Myc-tagged ubiquitin. Atg32-3×Strep-9×His was pulled down by using Ni-NTA Sepharose under denatured conditions in the presence of urea to dissociate proteins non-covalently bound to Atg32, and then was precipitated using Myc antibody-conjugated beads. Under these conditions, Atg32-3×Strep-9×His is still coprecipitated with Myc-tagged ubiquitin (Fig. 2D), suggesting that Atg32 is covalently bound to ubiquitin chains under respiratory conditions. However, we observed that Atg32-3×HA remained coimmunoprecipitated with ubiquitin in cells lacking Doa10 (Fig. S3A). Given that Atg32 stabilization was moderate in Doa10-deficient cells treated with CHX (Fig. 2B), these data raise the possibility that additional E3 ligases and/or other factors may mediate the destabilization of this mitophagy receptor in the absence of Doa10.

### Doa10 interacts with Atg32

To determine whether Doa10 recognizes Atg32, we tested Doa10–Atg32 interactions by Co-IP using cells expressing HA-tagged Atg32 and Myc-tagged Doa10. Since Doa10 substrates are supposed to be rapidly degraded via the UPS, Co-IP was performed using the cross-linker dithiobis (succinimidyl-propionate) (DSP) (Akaki et al, 2022). We found that Atg32-3×HA was coprecipitated with Doa10-9×Myc in the cross-linking Co-IP assay (Fig. 3A), indicating that Doa10 interacts with Atg32. This finding was further supported by the additional Co-IP assay using cells expressing Doa10^C39S^, where this catalytically inactive Doa10 mutant is also coprecipitated with Atg32-3×HA (Fig. 3B).

**Figure 3.**
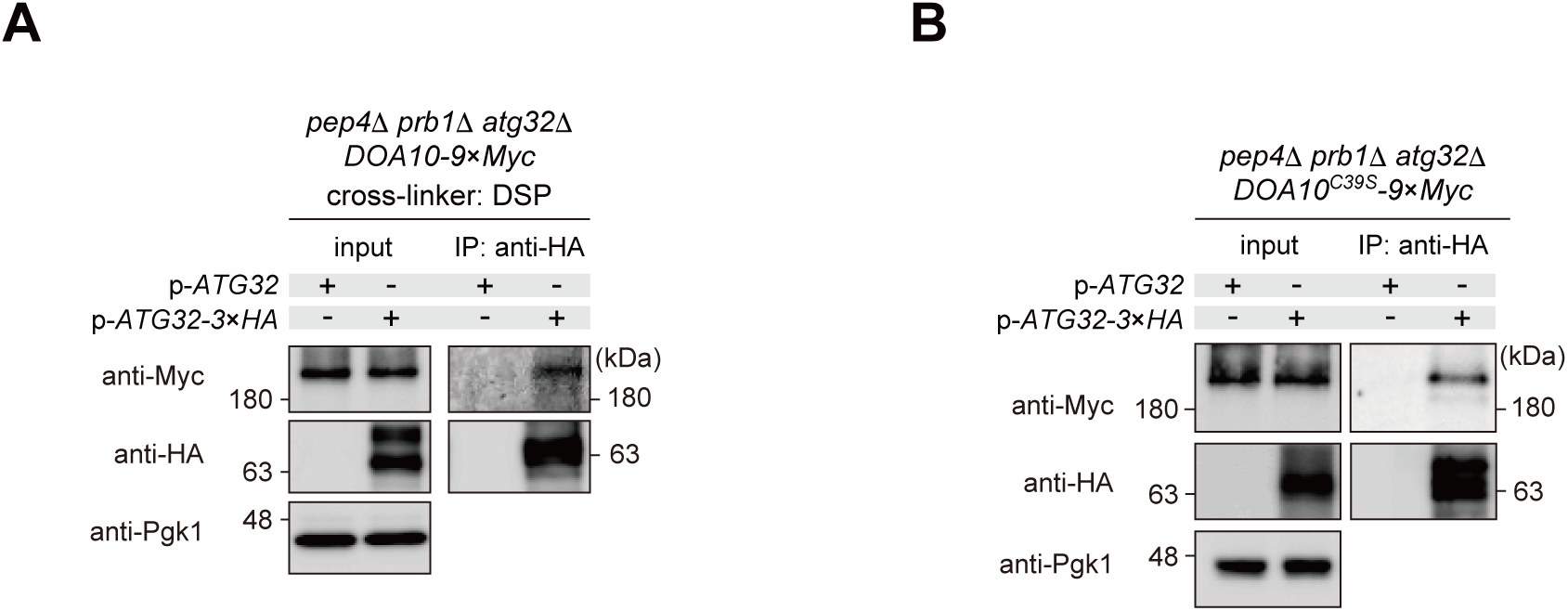
Doa10 interacts with Atg32. **(A)** *DOA10-9*×*Myc*-expressing *atg32*Δ cells harboring chromosome- or plasmid-encoded versions of Atg32 or Atg32-3×HA were grown in glycerol medium with the cross-linker DSP, and subjected to coimmunoprecipitation using anti-HA antibody-conjugated agarose. Eluted immunoprecipitates (IP) and detergent-solubilized mitochondria-enriched fractions (input) were analyzed by western blotting. Pgk1 was monitored as a loading control. **(B)** *DOA10^C39S^-9×Myc*-expressing *atg32*Δ cells harboring chromosome- or plasmid-encoded versions of Atg32 or Atg32-3*×*HA were grown in glycerol medium, and subjected to coimmunoprecipitation using anti-HA antibody-conjugated agarose. Eluted immunoprecipitates (IP) and detergent-solubilized mitochondria-enriched fractions (input) were analyzed by western blotting. Pgk1 was monitored as a loading control.

### The GET pathway is required for ER targeting of Atg32

Since the Doa10 complex is localized in the ER membrane, we hypothesized that Atg32 is targeted to the ER and recognized by Doa10. To test this hypothesis, we constructed cells expressing 3×GFP-tagged Atg32 from its endogenous promoter, allowing us to observe mitochondrial patterns of this protein (Fig. 4A). Notably, we observed that a fraction of Atg32 exhibits ER localization in *doa10*-null cells but not in wild-type cells (Fig. 4A and B), suggesting that Atg32 is targeted and accumulated on the ER upon loss of this E3 ligase.

**Figure 4.**
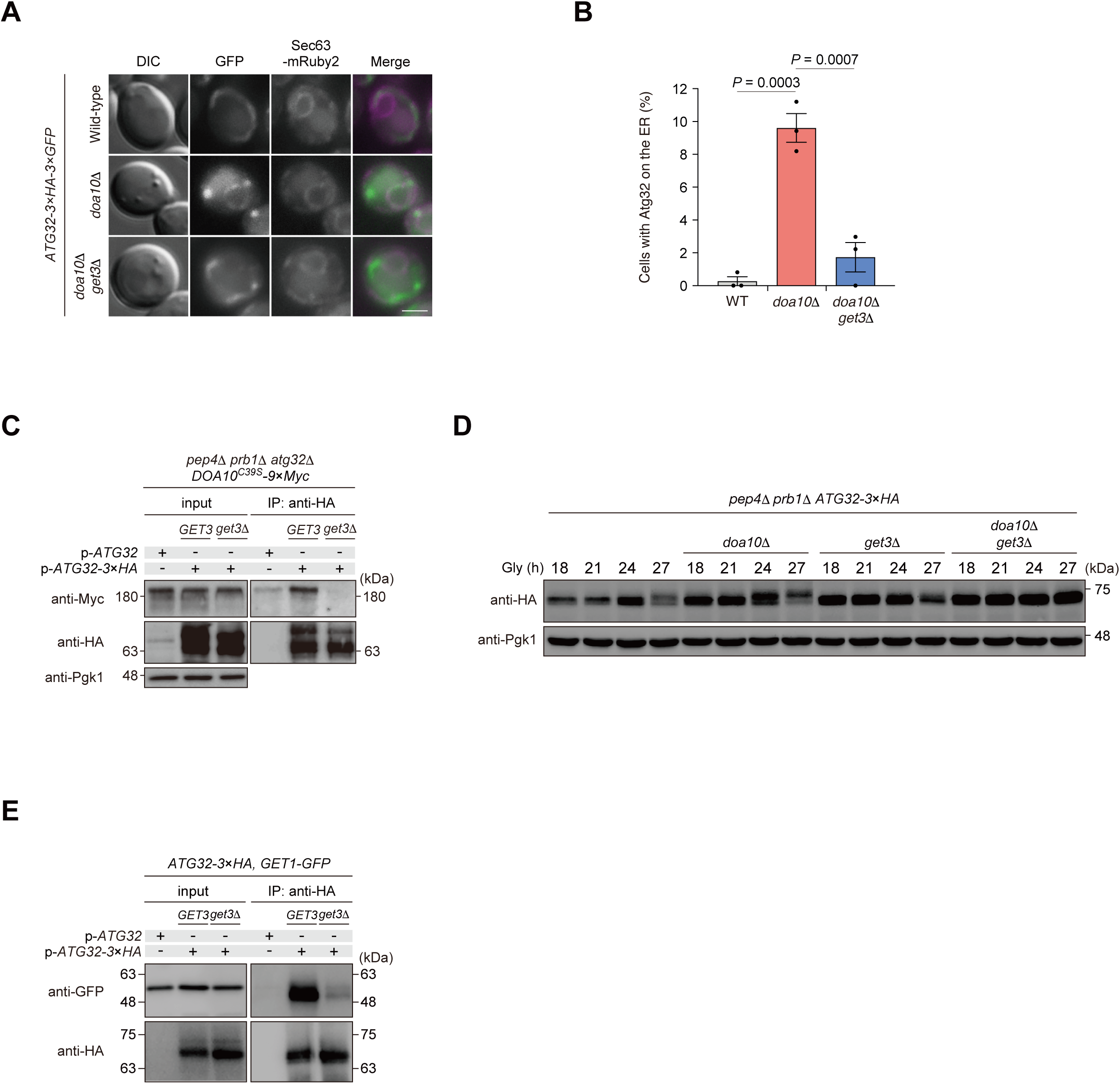
Atg32 is targeted to the ER in a manner dependent on the GET pathway. **(A)** Representative images of Atg32-3×HA-3×GFP and Sec63-mRuby2-expressing wild-type, *doa10*Δ cells and *doa10*Δ *get3*Δ cells grown for 24 h in dextrose medium. Single-plane images are shown. Scale bar, 2 μm. DIC, differential interference contrast. **(B)** Cells analyzed in **(A)** were quantified in three experiments. Data represent the averages of all experiments (*n* = 3 independent cultures, means ± s.e.m.). **(C)** *DOA10^C39S^-9×myc*-expressing wild-type and *get3*Δ cells harboring chromosome- or plasmid-encoded versions of Atg32 or Atg32-3*×*HA were grown in glycerol medium, and subjected to coimmunoprecipitation using anti-HA antibody-conjugated agarose. Eluted immunoprecipitates (IP) and detergent-solubilized mitochondria-enriched fractions (input) were analyzed by western blotting. Pgk1 was monitored as a loading control. **(D)** Atg32-3*×*HA-expressing wild-type, *doa10*Δ*, get3*Δ and *doa10*Δ *get3*Δ cells were grown in glycerol medium (Gly), collected at the indicated time points, and subjected to western blotting. Pgk1 was monitored as a loading control. **(E)** The endogenous Get1-GFP and Atg32-3*×*HA-expressing wild-type and *get3*Δ harboring chromosome- or plasmid-encoded versions of Atg32 or Atg32-3*×*HA were grown in glycerol medium, and subjected to coimmunoprecipitation using anti-HA antibody-conjugated agarose. Eluted immunoprecipitates (IP) and detergent-solubilized mitochondria-enriched fractions (input) were analyzed by western blotting. **(C, D)** All strains are *pep4 prb1*-double-null derivatives defective in vacuolar degradation of Atg32-3*×*HA via mitophagy. Data were analyzed by one-way ANOVA with Tukey’s multiple comparison test **(B)**.

A previous study has reported that non-imported mitochondrial carrier proteins are localized to the ER upon mitochondrial membrane potential dissipation dependently on the GET pathway (Vitali et al, 2018). This pathway is composed of a series of proteins coordinately acting to recognize the TA proteins as substrates, delivers them to the ER, and finally inserts them into the ER membrane. Among these GET components, the Get1/2 insertase complex on the ER receives TA proteins released from Get3, a cytosolic ATPase chaperone (Denic, 2012; Denic et al, 2013; Farkas & Bohnsack, 2021). To assess whether the GET pathway acts in the ER targeting of Atg32, we monitored localization of Atg32 in cells lacking Doa10 and Get3. In striking contrast, a number of cells containing ER-localized Atg32-3×GFP in *doa10*-null cells was significantly reduced in *doa10*- *get3*-null cells (Fig. 4A and B). Furthermore, ablation of Get3 impaired the Doa10^C39S^-Atg32 interaction (Fig. 4C). Together, these data suggest that the GET pathway targets Atg32 to the ER, which is required for Doa10-Atg32 interactions. Since Get3 seems to be required for the ER localization of Atg32 and its recognition by Doa10, we speculated that a fraction of Atg32 cannot be degraded and accumulated in *get3*-null cells. To test this possibility, we examined Atg32 protein levels in cells lacking Get3. Our western blot analysis showed that Atg32 protein levels were increased in the absence of Get3 under respiratory conditions (Fig. 4D), indicating that targeting of Atg32 to the ER via Get3 appears to be required for efficient protein turnover of Atg32.

To further test whether the Get1/2 insertase complex captures Atg32 as a substrate, we performed an immunoprecipitation assay using cells expressing Get1-GFP and Atg32-3×HA. Under respiratory conditions, Get1-GFP was coprecipitated with Atg32-3×HA, indicating that Get1/2 interacts with Atg32 (Fig. 4E). By contrast, in cells lacking Get3, the Get1–Atg32 interaction was significantly compromised (Fig. 4E), suggesting that Get3 is required for the Get1/2 complex to capture Atg32.

Recent studies have reported that Msp1, an N-terminally anchored mitochondrial AAA-ATPase, removes mistargeted proteins from the mitochondrial membrane, and reinserts them into the ER for degradation by Doa10 (Matsumoto et al, 2019; Dederer et al, 2019; Matsumoto et al, 2022). To examine whether Msp1 is involved in Atg32-mediated mitophagy, we examined mitophagy in *msp1*-null cells. Mitochondrial degradation in cells lacking Msp1 was almost comparable to that in wild-type cells (Fig. S4A and B), indicating that Msp1-mediated protein extraction from mitochondrial membranes is unlikely to be involved in the regulation of Atg32 and mitophagy.

### Increased hydrophobicity of the Atg32 transmembrane domain enhances the ER localization of Atg32

To gain insight into how the GET pathway captures Atg32, we focused on previous studies showing that ER membrane proteins have relatively high hydrophobicity in their transmembrane domains (TMDs), and that the GET pathway efficiently captures TA proteins with relatively highly hydrophobic TMDs (Wang et al, 2011; Rao et al, 2016; Guna et al, 2018). We therefore decided to investigate whether the hydrophobicity of the Atg32 TMD may determine its ER localization. To this end, we generated plasmids encoding 3×GFP-tagged Atg32 variants in which amino acid residues in the conserved TMDs (W390, W393, S396, G400 and G404) were substituted with alanine (Fig. 5A). The hydrophobicity of these amino acid residues was estimated from the change in free energy (*ΔG*) upon transition from the lipid to the aqueous layer (*ΔG* of each amino acid is S(0.6) < G(1.0) < A(1.6) < W(1.9)) (Engelman et al, 1986). Atg32 mutants with relatively reduced TMD hydrophobicity compared to wild-type (Atg32^W390A^ and Atg32^W393A^) showed mitochondrial patterns similar to those of wild-type Atg32, whereas Atg32 mutants with relatively increased TMD hydrophobicity (Atg32^S396A^, Atg32^G400A^ and Atg32^G404A^) were localized not only to mitochondria but also to the ER (Fig. 5B and C). The ER pattern of these Atg32 mutants (Atg32^S396A^, Atg32^G400A^ and Atg32^G404A^) is less significantly increased in cells deficient for Get3, which may suggest that increased hydrophobicity in the Atg32 TMD is associated with the enhanced ER targeting of Atg32 via Get3 (Fig. S4C and D). We also found that loss of Doa10 led to a much larger increase in the ER localization of these Atg32 mutants (Fig. 5B and C). excluding the possibility that they are defective in the interaction with Doa10. Conversely, loss of Doa10 fails to efficiently accumulate Atg32^W390A^ and Atg32^W393A^ mutants with relatively reduced TMD hydrophobicity to the ER (Fig. 5D and E), suggesting that the GET pathway may not efficiently capture and target Atg32 mutants with a less hydrophobic TMD to the ER.

**Figure 5.**
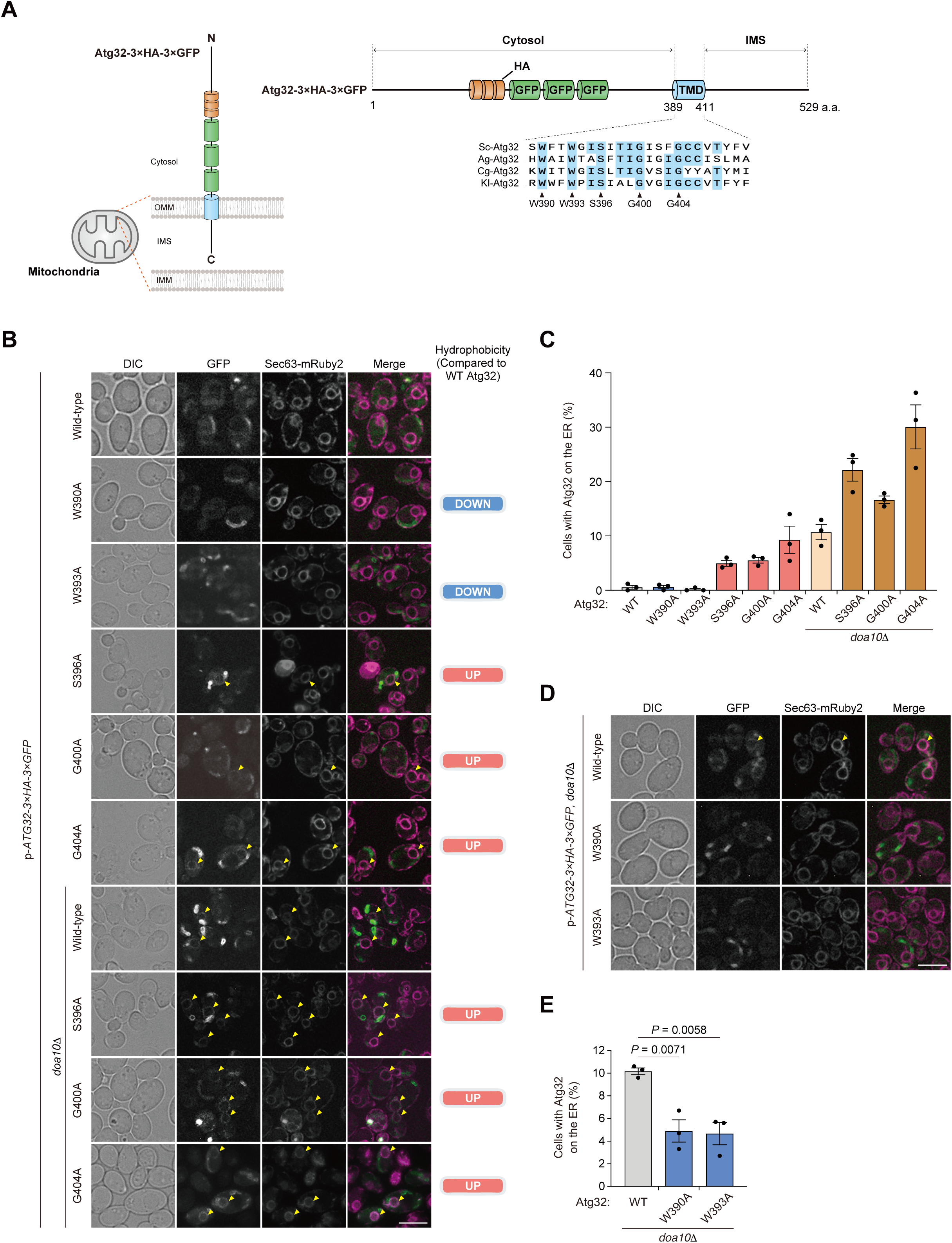
Increased hydrophobicity of the Atg32 transmembrane domain enhances the ER localization of Atg32. **(A)** Schematic illustration of 3×HA- and 3×GFP-tagged Atg32 topology with the transmembrane domain (TMD) and the cytosolic domains. The amino acid region of *Saccharomyces cerevisiae* Atg32 aligned using Clustal W2 with Atg32 homologs from *Sc*, *S. cerevisiae*; *Ag*, *Ashbya gossypii*; *Cg*, *Candida glabrata*; *Kl*, *Kluyveromyces lactis*. **(B)** Representative structured illumination microscopy images of wild-type and *doa10*Δ cells expressing Sec63-mRuby2 harboring plasmid-encoded versions of Atg32-3×HA-3×GFP were grown in dextrose medium (SDCA) to OD_600_ = 1.6, and observed under a fluorescence microscopy. Yellow arrowheads indicate Atg32-3×HA-3×GFP overlapping Sec63-mRuby2. Scale bar, 5 μm. DIC, differential interference contrast. **(C)** Cells with Atg32-3×HA-3×GFP on the ER in the images were quantified in three experiments. Data represent the averages of all experiments (*n* = 3 independent cultures, means ± s.e.m.). **(D)** Representative structured illumination microscopy images of wild-type and *doa10*Δ cells expressing Sec63-mRuby2 harboring plasmid-encoded versions of Atg32-3×HA-3×GFP grown in dextrose medium (SDCA) to OD_600_ = 1.6, and observed using fluorescence microscopy. Yellow arrowheads indicate Atg32-3×HA-3×GFP overlapping Sec63-mRuby2. Scale bar, 5 μm. DIC, differential interference contrast. **(E)** Cells with Atg32-3×HA-3×GFP on the ER in the images were quantified in three experiments. Data represent the averages of all experiments (*n* = 3 independent cultures, means ± s.e.m.). Data were analyzed by one-way ANOVA with Dunnett’s multiple comparison test **(E)**.

### A region adjacent to the transmembrane domain is required for Doa10–Atg32 interactions

To define the region of Atg32 required for recognition by Doa10, we expressed a series of 3×GFP-tagged C-terminal truncation mutants under the endogenous promoter and examined their ER localizations (Fig. 6A). We found that Atg32(1–529), Atg32(1–508), and Atg32(1–488) rarely accumulated on the ER in wild-type cells, whereas ER localization was markedly increased in *doa10*-null cells (Fig. 6B and C; Fig. S5A). In contrast, further truncation to the residue 468 resulted in an increase in ER localization even in the presence of Doa10 (19%). Atg32(1–468) was accumulated on the ER at 2-fold higher levels compared to cells lacking Doa10, suggesting that the residues 469–488 are required for proper recognition of Atg32 by Doa10. A similar localization pattern was observed for Atg32(1–448) (19%), whereas additional truncation to residue 418 partially reduced ER accumulation (12%). These observations prompted us to test whether this region contributes to the interaction of Atg32 with Doa10. To this end, we performed co-immunoprecipitation assays using the catalytically inactive Doa10^C39S^ mutant. Consistent with our previous results, full-length Atg32 efficiently coprecipitated with Doa10^C39S^ (Fig. 6D, Lane No.3). In contrast, Atg32(1–468) mostly failed to coprecipitate with Doa10^C39S^ despite being recovered at comparable levels in the immunoprecipitated fraction (Fig. 6D, Lane No.2). These data are consistent with the idea that the C-terminal region downstream of residue 468 is required for the efficient interaction between Atg32 and Doa10.

**Figure 6.**
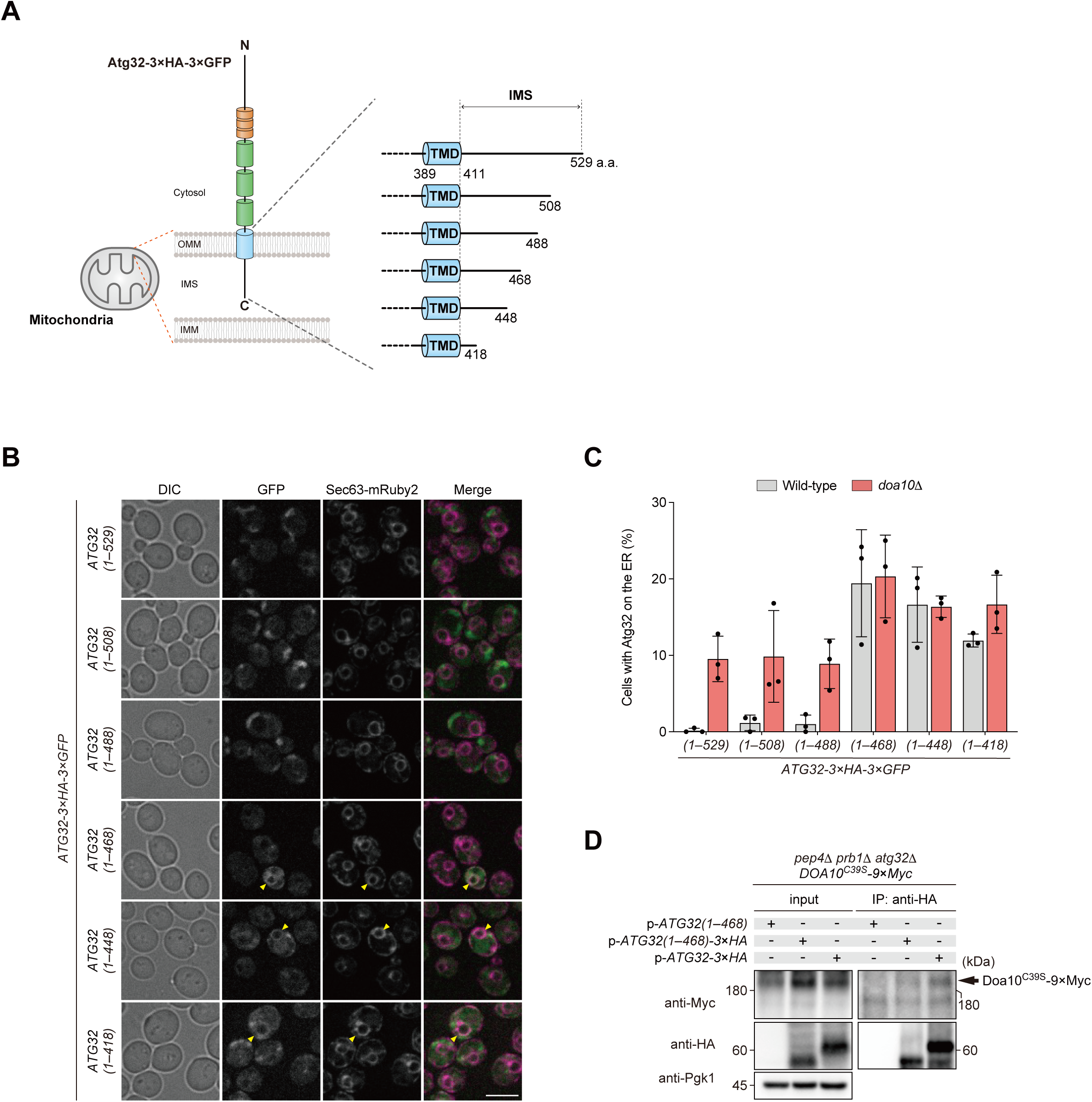
A region adjacent to the transmembrane domain is required for Doa10–Atg32 interactions. **(A)** Schematic illustration of full-length and C-terminally truncated 3×HA- and 3×GFP-tagged Atg32 constructs. A series of C-terminal truncation mutants terminating at amino acid residues 508, 488, 468, 448, and 418 were generated from the full-length protein (529 a.a.). **(B)** Representative structured illumination microscopy images of Sec63-mRuby2-expressing cells chromosomally expressing truncated variants of Atg32-3×HA-3×GFP grown in dextrose medium (SDCA) to OD_600_ = 1.6. Yellow arrowheads indicate Atg32-3×HA-3×GFP overlapping with Sec63-mRuby2. Scale bar, 5 μm. DIC, differential interference contrast. **(C)** Cells with Atg32-3×HA-3×GFP on the ER in the presence (wild-type) or absence (*doa10*1−) of Doa10 were quantified in three experiments. Data represent the averages of all experiments (*n* = 3 independent cultures, means ± s.e.m.). **(D)** *DOA10^C39S^-9×myc*-expressing cells harboring plasmid-encoded versions of Atg32-3*×*HA, Atg32(1–468), and Atg32(1–468)-3*×*HA were grown in glycerol medium and subjected to coimmunoprecipitation using anti-HA antibody-conjugated agarose. Eluted immunoprecipitates (IP) and detergent-solubilized mitochondria-enriched fractions (input) were analyzed by western blotting. Pgk1 was monitored as a loading control. These strains are *pep4 prb1*-double-null derivatives defective in vacuolar degradation of truncated variants of Atg32-3*×*HA via mitophagy.

### ER degradation is enhanced in Doa10-deficient cells that overaccumulate Atg32

A previous study suggests that, since the cytosol domain of Atg32 contains conserved binding motifs to Atg8 (a ubiqutin-like protein localized to the autophagosome) and Atg11 required for localizing autophagosomes to mitochondria, the peroxisome-anchored Atg32 cytosolic domain acts as an autophagic degron targeting peroxisomes and mediating pexophagy (Kondo-Okamoto et al, 2012). This finding raises the possibility that faithful mitochondrial targeting of Atg32 is crucial for protecting other organelles from dysregulated or unnecessary degradation. We thus expected that over-accumulation and over-activation of Atg32 on the ER in the absence of Doa10 can increase ER degradation via autophagy (ER-phagy). To test this hypothesis, we first generated cells overexpressing Atg32 under a TEF promoter. To clearly see the ER-phagic degradation potentially induced by Atg32, we knocked out Atg39 and Atg40 to inhibit the conventional ER-phagy pathway. To maximize the activity of Atg32 to initiate autophagic degradation, we also deleted Ppg1, a negative regulator of Atg32 that is dually localized to the mitochondrial surface and the ER (Furukawa et al, 2018; Innokentev et al, 2020; Onishi et al, 2023). In *atg39*- and *atg40*-double null cells (conventional ER-phagy-deficient cells), the absence of Doa10 increases the processing of Sec63-mCherry when Ppg1 is deleted and Atg32 is over-accumulated (Lane No. 12 and 16 in Fig. 7A and B). This implicates that increased accumulation of Atg32 on the ER upon loss of Doa10 may result in degradation of the ER.

**Figure 7.**
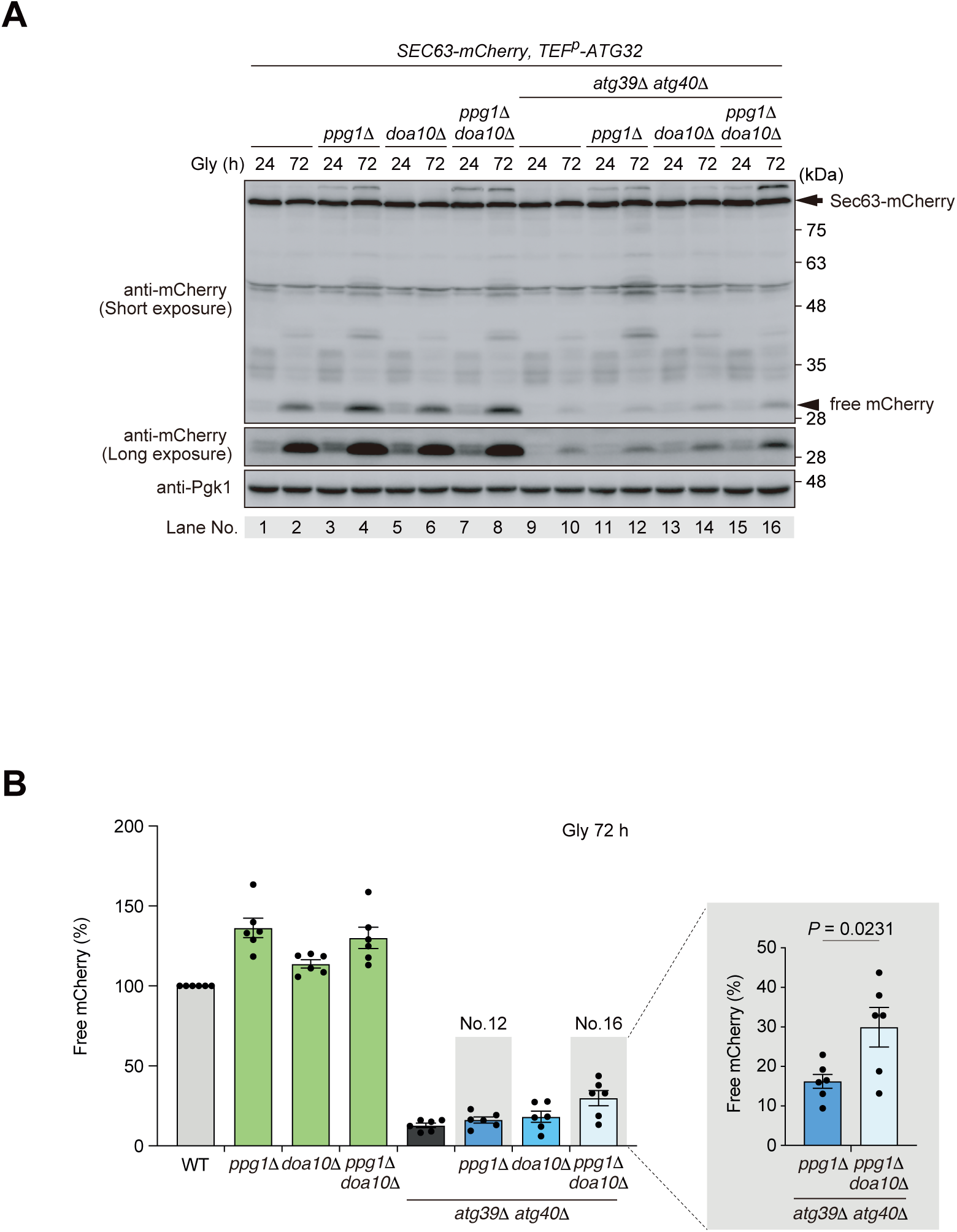
ER-phagy is accelerated in cells lacking Doa10 and overaccumulating Atg32. **(A)** Sec63-mCherry-expressing wild-type, *ppg1*Δ*, doa10*Δ, *ppg1*Δ *doa10*Δ, *atg39*Δ *atg40*Δ, *atg39*Δ *atg40*Δ *ppg1*Δ, *doa10*Δ *atg39*Δ *atg40*Δ, and *ppg1*Δ *doa10*Δ *atg39*Δ *atg40*Δ cells were grown in glycerol medium (Gly), collected at the indicated timepoints, and subjected to western blotting. Generation of free mCherry indicates transport of the maker to the vacuole. Pgk1 was monitored as a loading control. **(B)** The amounts of free mCherry in cells analyzed in **(A)** were quantified in six experiments. The signal intensity value of free mCherry in wild-type cells at the 72 h time point was set to 100%. Data represent the averages of all experiments (*n* = 6 independent cultures, means ± s.e.m.). Data were analyzed by a two-tailed *t* test **(B)**.

## DISCUSSION

Our results demonstrate that the ERAD Doa10 E3 ligase acts to downregulate mitophagy by mediating degradation of functional Atg32 molecules (Fig. 8A and B), adding this mitophagy receptor to the list of potential endogenous ERAD substrates (Foresti et al, 2013; Foresti et al, 2014). This degradation of Atg32 seems to be mediated by the UPS, as Atg32 is covalently linked to ubiquitin chains and its degradation is partially suppressed by the MG132 treatment (Fig. 2A, C, and D). These results are consistent with a previous study where the Atg32 ubiquitination site was determined and expression of a ubiquitin-deficient Atg32 mutant increases mitophagy (Camougrand et al, 2020). We also show that the ER is a site where a fraction of Atg32 is localized and recognized by the Doa10 E3 ligase, which appears to be mediated by the GET pathway (Fig. 4A, B, C and D). This is in agreement with the requirement of the GET pathway for targeting mitochondrial proteins to the ER under certain conditions such as mitochondrial membrane depolarization (Vitali et al, 2018; Xiao et al, 2021; Matsumoto et al, 2022). Collectively, we suggest that the ERAD and GET pathways may act in concert with each other to reduce Atg32 protein levels. Consistently, the ablation of the Doa10 E3 ligase increases Atg32 protein levels and enhances mitophagic degradation (Fig. 1D and E).

**Figure 8.**
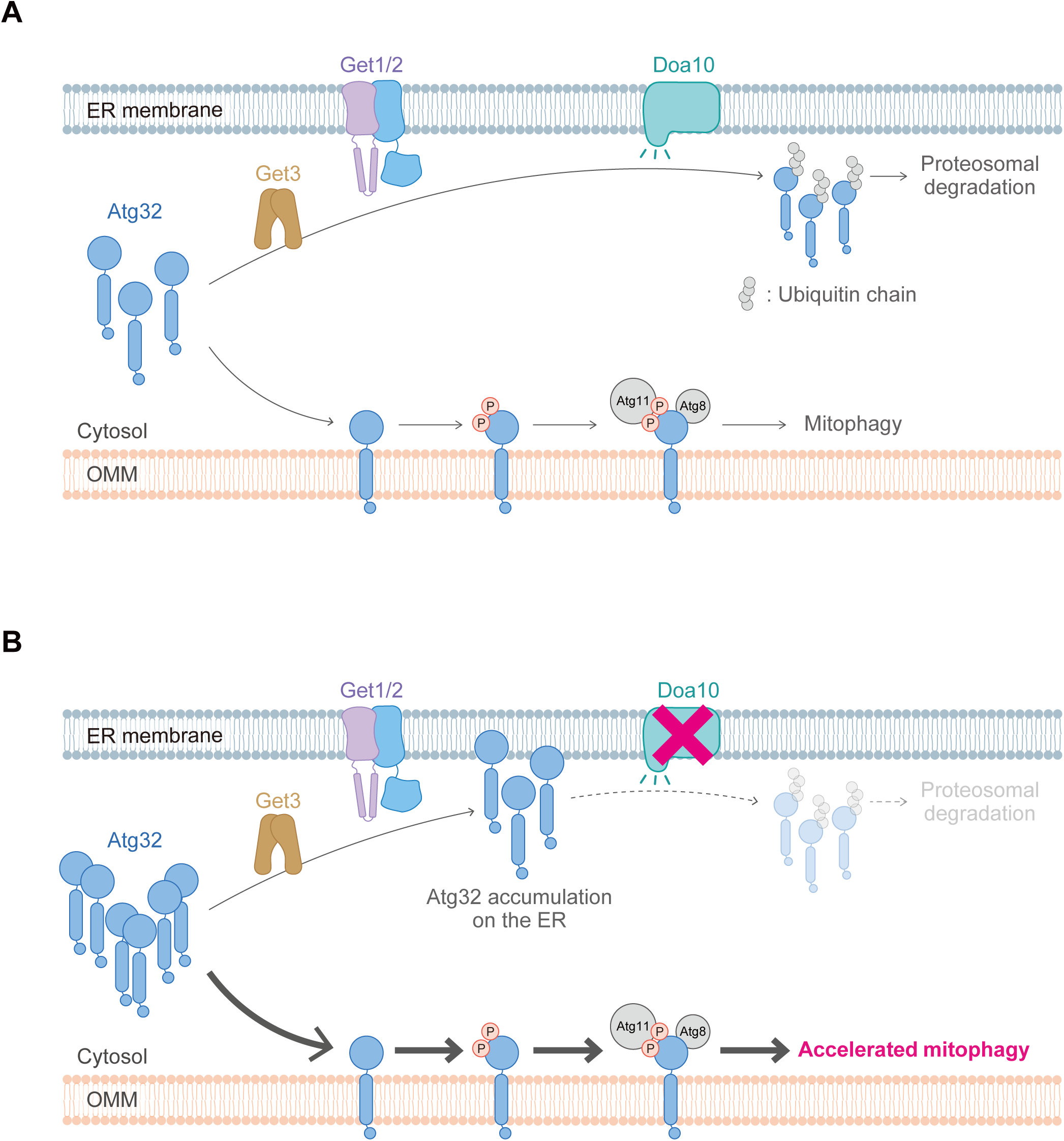
A working model for the GET- and ERAD-mediated brake control of Atg32-dependent mitophagy. **(A)** Shortly after the release from the ribosome, a fraction of Atg32 is captured by the GET pathway, and targeted to the ER. The Doa10 E3 ligase complex recognizes and ubiquitinates Atg32 for its degradation by the proteasome. **(B)** Deficiency of Doa10 retards Atg32 protein turnover and accumulates Atg32 on the ER, leading to increased Atg32 protein levels and promoting mitophagy.

Our previous results showed that the GET pathway-dependent ER targeting of the Ppg1 complex, a multi-subunit phosphatase complex that suppresses Atg32 phosphorylation and mitophagy, is important to prevent dysregulated Atg32 phosphorylation and support mitophagy activation (Onishi et al, 2023). On the other hand, we propose here that the GET pathway appears to target Atg32 to the ER, allowing Doa10 to recognize this mitophagy receptor and ultimately limit mitophagy (Fig. 8A and B). This leads us to speculate the possibility that the GET pathway may capture different membrane proteins and target them to the ER, fine-tuning Atg32 activity by modulating its post-translational modification (phosphorylation) and its protein levels. It remains to be determined whether the GET pathway controls the membrane insertion of different clients into the ER in a context- or time-dependent manner upon mitophagy induction, to further provide a regulatory axis to modulate this catabolic pathway.

The mechanisms by which mitochondrial proteins, including Atg32, are targeted to the ER are not fully understood. Although the GET pathway is initiated shortly after release of TA proteins from ribosomes (Zhang et al, 2021), it remains possible that mitochondrial membrane proteins are redirected to the ER after being localized on the mitochondrial surface. Accumulating evidence suggests that Msp1 extracts inappropriately targeted non-mitochondrial TA proteins from mitochondrial membranes (Chen et al, 2014; Okreglak & Walter, 2014; Wohlever et al, 2017; Wang et al, 2020; Matsumoto et al, 2022). We therefore speculated that Msp1 might act in the extraction of Atg32 and its redirection to the ER, however, we did not observe significant changes in mitophagy levels in cells lacking Msp1 (Fig. S4A and B). Taken together, it seems unlikely that Msp1-mediated protein extraction from the OMM is involved in Atg32-mediated mitophagy. Rather, the GET pathway may directly capture Atg32 prior to its insertion into the OMM. Consistent with this idea, deletion of the residues 469–488 markedly increased ER localization of the Atg32(1–468) truncation mutant even in the absence of Doa10 (Fig. 6B and C). Thus, the elevated ER localization of Atg32(1–468) may not simply reflect its impaired interaction with Doa10 on the ER, but instead also suggest that this region bifunctionally contributes to efficient mitochondrial targeting of Atg32. Our possible explanation for these phenotypes is that loss of this region could increase the fraction of newly synthesized Atg32 mistargeted to the ER before its insertion into the OMM. Future studies will be required to determine whether the residues 469–488 participate directly in mitochondrial targeting of Atg32 and recognition by the GET pathway.

We demonstrated that Atg32 is recognized by the Doa10 E3 ligase and covalently attached to ubiquitin chains, however, Atg32 stabilization upon loss of Doa10 was moderate (Fig. 2B). In addition, we still detected ubiquitin signals coprecipitated with Atg32 in the absence of Doa10 (Fig. S3A), together raising the possibility that additional E3 ligases and/or proteases may mediate the degradation of Atg32. MARCH5, a mitochondrial E3 ligase, has been reported to ubiquitinate FUNDC1, a mitophagy receptor to fine-tune hypoxia-induced mitophagy in mammalian cells (Liu et al, 2012; Chen et al, 2017). Also, the SCF-FBXL4 ubiquitin E3 ligase complex is shown to ubiquitinate the NIX and BNIP3 mitophagy receptors to limit basal mitophagy in mammals (Nguyen-Dien et al, 2023; Cao et al, 2023; Elcocks et al, 2023). In addition, Yme1, a catalytic subunit of the mitochondrial inner membrane AAA protease, appears to proteolytically process Atg32 (Wang et al, 2013). Thus, it remains plausible that multiple E3 ligases/proteases act in a redundant manner to degrade Atg32.

Targeting of Atg32 to the other organelles, such as peroxisomes, can lead to autophagic degradation of these organelles (Kondo-Okamoto et al, 2012). The absence of Doa10 increases ER degradation, presumably via autophagy, in Ppg1-, Atg39-, and Atg40-deficient cells overexpressing Atg32 (Fig. 7A and B), suggesting that the Doa10-mediated degradation of this pro-mitophagic protein may prevent dysregulated degradation of the ER. A previous study identified the ER-resident P5A-ATPase ATP13A1 (Spf1 in yeast) that extracts mistargeted mitochondrial TA proteins from the ER membrane (McKenna et al, 2020; McKenna et al, 2022). It is thus possible that in the absence of Doa10, the Spf1 extractase may act in removing Atg32 localized at the ER surface. Whether and how the GET-dependent insertion, ERAD-mediated protein turnover, and the extractase-driven removal of membrane proteins coordinately act to regulate Atg32 and mitochondrial degradation in a spatio-temporal manner remains to be elucidated in future studies.

## MATERIALS & METHODS

### Growth conditions of yeast

Yeast cells were incubated in YPD medium (1% yeast extract, 2% peptone, and 2% dextrose), synthetic medium (0.17% yeast nitrogen base without amino acids and ammonium sulfate, 0.5% ammonium sulfate) with 0.5% casamino acids containing 2% dextrose (SDCA), or 0.1% dextrose plus 3% glycerol (SDGlyCA), supplemented with necessary amino acids. For induction of Atg32 and mitophagy under respiratory conditions, cells grown to mid-log phase in SDCA were transferred to SDGlyCA and incubated at 30 °C.

### Western blotting

Samples corresponding to 0.1 OD_600_ units of cells were separated by SDS-PAGE followed by western blotting and immunodecoration with primary antibodies raised against mCherry (1:2,000, Abcam ab125096), Pgk1 (1:10,000, Abcam, ab113687), GFP (1:1,000, Roche, 13921700), HA (1:5,000, Sigma, A2095), anti-Ub, anti-Strep, and anti-Myc. After treatment with the secondary antibodies, horseradish peroxidase (HRP)-conjugated rabbit anti-mouse IgG (H + L) for mCherry, GFP, HA, Pgk1, followed by the enhanced chemiluminescence reagent Western Lightning Plus-ECL (PerkinElmer, 203-19151) or ImmunostarLD (Wako, PTJ2005). After treatment with enhanced chemiluminescence reagents, proteins were detected using a luminescence image analyzer (LAS-4000 mini; GE Healthcare) or a luminescence image analyzer (FUSION Solo S; VILBER). Quantification of the signals was performed using ImageQuant TL (GE Healthcare).

### Treatment with the proteosome inhibitor

MG132 (50 µM, SI9710; LifeSensors) was added to mid-log phase cell culture in SDGlyCA. Samples were prepared from 1.0 OD_600_ units of cells collected at the indicated time points and analyzed by western blotting.

### Immunoprecipitation

Coimmunoprecipitation assays were performed using a vacuolar protease-deficient strain transformed with a plasmid encoding Atg32 or Atg32-3×HA. 90–180 OD_600_ units of cells grown in SDGlyCA for 24 h were collected by centrifugation, washed once with H2O, resuspended in TD buffer (0.1 M Tris-SO_4_ [pH 9.4], 10 mM DTT), and incubated for 10 min at 30 °C. Cells were collected by centrifugation, resuspended in SP buffer (20 mM potassium phosphate buffer [pH 7.4], 1.2 M sorbitol) containing 200 µg/ml zymolyase 100T (120493; Seikagaku), and incubated for 60 min at 30 °C. Spheroplasts were washed with SP buffer and resuspended in SH buffer (0.6 M sorbitol, 20 mM HEPES-KOH [pH 7.4] containing protease inhibitor mixture (1861284; Thermo Scientific). Whole cell homogenates were subjected to centrifugation (500 × g) at 4 °C for 5 min. Membrane and soluble fractions were separated by centrifugation (17,000 × g) at 4 °C for 5 min. Mitochondria-enriched fractions were then resuspended in lysis buffer (50mM Tris-HCl [pH 7.5], 100 mM NaCl, 0.1 mM EDTA, 0.4% Triton X-100, and protease inhibitor mixture) at 4 °C for 8 min and subjected to centrifugation (17,000 × g) at 4 °C for 5 min. The supernatant was incubated with 30 µl of anti-HA-agarose conjugate (A2095; Sigma) at 4 °C for 2 h with gentle agitation. The beads were washed twice with wash buffer (50 mM Tris-HCl [pH 7.5], 300 mM NaCl, 0.1 mM EDTA, 0.4% Triton X-100, and protease inhibitor mixture), and once with PBS. Immunoprecipitates were eluted with SDS sample buffer and analyzed by western blotting.

In the case of cross-linking, we followed the previously reported method (Gardner et al, 2000) and modified our immunoprecipitation method. After incubation of cells in SP buffer containing zymolyase for 30 min at 30 °C, dithiobis (succinimidyl-propionate) (DSP) (D629; DOJINDO) was added at a final concentration of 400 µg/ml. Cells were incubated for 30 min at 30 °C. Tris (200 mM, pH 7.5) was added to quench the cross-linking reaction. Spheroplasts were incubated for 10 min at 30 °C and washed twice with SP buffer, then procedures were performed as described above.

### Tandem protein purification (TAP)

TAP assays were performed using an Atg32-3×Strep-9×His expressing vacuolar protease-deficient strain transformed with a plasmid encoding Ub or Ub-Myc. Mitochondria-enriched fractions were obtained from the 1500–1800 OD_600_ units of cells grown in SDGlyCA for 24 h as described in the immunoprecipitation method. Mitochondria-enriched fractions were resuspended in lysis buffer (50 mM Tris-HCl [pH 7.5], 100 mM NaCl, 0.1 mM EDTA, 0.4% Triton X-100, and 1 mM Phenylmethylsulfonyl fluoride (PMSF)) at 4 °C for 30 min and subjected to centrifugation (17,000 × g) at 4 °C for 5 min. Urea (6M) were added, and then the supernatant was incubated with 500 µl of Ni-NTA agarose (R90101; Thermo Scientific) at 4 °C for overnight with gentle agitation in a denatured condition. The agarose was washed once with wash Imidazole buffer (50 mM Tris-HCl [pH 7.5], 300 mM NaCl, 0.1 mM EDTA, 0.4% Triton X-100, 20 mM Imidazole (pH7.5), and protease inhibitor mixture), and twice with wash buffer. Pull-down samples were eluted with Ni-NTA elution buffer (50 mM Tris-HCl [pH 7.5], 100 mM NaCl, 0.1 mM EDTA, 0.4% Triton X-100, 250 mM Imidazole (pH7.5)). The eluted solution was incubated with 30 µl of anti-Myc-agarose conjugate (A7470; Sigma) at 4 °C for 2 h with gentle agitation. The beads were washed three times with PBS. Immunoprecipitates were eluted with SDS-sample buffer and analyzed by western blotting.

### Cycloheximide chase assay

Cycloheximide (200 µg/ml, 033-20993; Wako) was added to mid-log phase cell culture in SDGlyCA. Samples were prepared from 1.0 OD_600_ units of cells collected at the indicated time points and analyzed by western blotting.

### Fluorescence microscopy

Live yeast cells expressing Atg32-3×GFP in Fig. 4A were observed using an inverted microscope (Axio Observer. Z1; Carl Zeiss) equipped with differential interference contrast optics, epifluorescence capabilities, a 100× objective lens (αPlan-APOCHROMAT 100, NA: 1.46; Carl Zeiss), a monochrome CCD camera (AxioCam MRm; Carl Zeiss), and filter sets for green fluorescent protein (GFP) derived from the jellyfish Aequorea victoria and mCherry, a monomeric version of the red fluorescent protein DsRed derived from the reef corals Discosoma sp. (13 and 20, respectively; Carl Zeiss). Cell images were captured using acquisition and analysis software (Axio Vision 4.6; Carl Zeiss).

### Structured illumination microscopy

Live yeast cells expressing Atg32-3×GFP in Fig. 5B and D were observed using structured illumination microscopy. Differential interference contrast and fluorescence images were obtained under a KEYENCE BZ-X810 system equipped with a 100× objective lens (CFI Apochromat TIRF 100XC Oil, Plan-APO TIRF 100, NA: 1.49; Nikon), filter sets for GFP and mCherry (BZ-X filter GFP and BZ-X filter TRITC, respectively; KEYENCE). Cell images were captured using acquisition and analysis software (BZ-X800 Analyzer; KEYENCE).

### Statistical analysis

Results are presented as means including means ± s.e.m. Statistical analyses were performed with Excel for Mac (Microsoft) and GraphPad Prism 9 (GraphPad Software), using a two-tailed *t*-test and one-way ANOVA followed by Tukey’s or Dunnett’s multiple comparison test. All the statistical tests performed are indicated in the figure legends.

## Acknowledgments

We thank Elmar Schiebel (Heidelberg University, Germany) for kindly providing us with the plasmid pFA6a-3myeGFP-kanMX6. This work was supported in part by JSPS KAKENHI Grants JP21K15041 (to M.O.), and JP20H05324 (to KO).

