## Supplemental figures for "The ERAD ubiquitin ligase Doa10 negatively regulates Atg32-mediated mitophagy"

**Short title:** The Doa10 complex reduces Atg32 protein levels and limits mitophagy

**Keywords:** ERAD, Doa10, GET pathway, Atg32, mitochondria, mitophagy

### SUPPLEMENTAL FIGURE LEGENDS

Figure S1. **Overexpression of Atg32 increases mitophagic degradation. (A)** Atg32-3×HA- and mito-DHFR-mCherry-expressing wild-type cells transformed with empty vectors or a plasmid encoding Atg32–3×HA were grown for the indicated time points in glycerol medium (Gly) and subjected to western blotting. Generation of free mCherry indicates transport of the maker to the vacuole. Pgk1 was monitored as a loading control. **(B)** The amounts of free mCherry in cells analyzed in **(A)** were quantified in three experiments. The signal intensity value of free mCherry in wild-type cells at the 24 h time point was set to 100%. Data represent the averages of all experiments ($n$ = 3 independent cultures, means ± s.e.m.). Data were analyzed by a two-tailed $t$ test **(B)**.

Figure S2. **Effects of loss of Doa10 on other types of selective autophagy and bulk autophagy. (A)** Wild-type, *doa10*Δ, *atg7*Δ cells expressing Tdh3-mCherry (a cytosol marker) were grown in glycerol medium (Gly), collected at the indicated time points, and subjected to western blotting. Generation of free mCherry indicates progression of bulk autophagy. Atg7 is an E1 enzyme essential for autophagy. The amounts of free mCherry in cells under respiratory conditions for 24 h, 48 h, and 72 h were quantified in three experiments. **(B)** The amounts of free mCherry in cells analyzed in **(A)** were quantified in three experiments. The signal intensity value of free mCherry in wild-type cells at the 72 h time point was set to 100%. Data represent the averages of all experiments ($n$ = 3 independent cultures, means ± s.e.m.). **(C)** Wild-type, *doa10*Δ, and *atg19*Δ cells expressing mito-DHFR-mCherry were grown in glycerol medium (Gly), collected at the indicated time points, and subjected to western blotting. Precursor form of Ape1 is transported from the cytosol to the vacuole via the Cvt pathway, a selective autophagy-related process, and cleaved to be a mature form. Atg19 is an adaptor protein required for the Cvt pathway. **(D)** Wild-type, *doa10*Δ, and *atg36*Δ cells expressing Pot1-mCherry (a peroxisome marker) were grown in glycerol medium (Gly), collected at the indicated time points, and subjected to western blotting. Generation of free mCherry indicated transport of peroxisomes to the vacuole. Atg36 is required for pexophagy. **(E)** The amounts of free mCherry in cells analyzed in **(D)** were quantified in three experiments. The signal intensity value of free mCherry in wild-type cells at the 72 h time point was set to 100%. Data represent the averages of all experiments ($n$ = 3 independent cultures, means ±

s.e.m.). **(F)** Wild-type, *doa10*Δ, and *atg39*Δ *atg40*Δ cells expressing Sec63-mCherry (an ER marker) were grown in glycerol medium (Gly), collected at the indicated time points, and subjected to western blotting. Generation of free mCherry indicated transport of ER to the vacuole. Atg39 and Atg40 are required for ER-phagy. **(G)** The amounts of free mCherry in cells analyzed in **(F)** were quantified in three experiments. The signal intensity value of free mCherry in wild-type cells at the 48 h time point was set to 100%. Data represent the averages of all experiments (*n* = 3 independent cultures, means ± s.e.m.).

Figure S3. **Ubiquitin signals are coprecipitated with Atg32 in the absence of Doa10. (A)** Myc-tagged ubiquitin-expressing *atg32*Δ or *atg32*Δ *doa10*Δ cells harboring plasmid-encoded versions of Atg32 or Atg32-3×HA were grown in glycerol medium and subjected to coimmunoprecipitation using anti-HA antibody-conjugated agarose. Eluted immunoprecipitates (IP) and detergent-solubilized mitochondria-enriched fractions (input) were analyzed by western blotting. Pgk1 was monitored as a loading control.

Figure S4. **Loss of Msp1 does not significantly affect mitophagy. (A)** Mitochondria-targeted DHFR-mCherry-expressing (mito-DHFR-mCherry) wild-type, *doa10*Δ, *msp1*Δ, and *atg32*Δ cells were grown for the indicated time points in glycerol medium (Gly), and subjected to western blotting. Generation of free mCherry indicates transport of the maker to the vacuole. Pgk1 was monitored as a loading control. **(B)** The amounts of free mCherry in cells analyzed in **(A)** were quantified in three experiments. The signal intensity value of free mCherry in wild-type cells at the 30 h time point was set to 100%. Data represent the averages of all experiments (*n* = 3 independent cultures, means ± s.e.m.). **(C)** Representative structured illumination microscopy images of *get3*Δ cells expressing Sec63-mRuby2 harboring plasmid-encoded versions of Atg32-3×HA-3×GFP were grown in dextrose medium (SDCA) to $OD_{600}$ = 1.6, and observed under a fluorescence microscopy. Scale bar, 5 μm. DIC, differential interference contrast. **(D)** Cells with Atg32-3×HA-3×GFP on the ER in the images were quantified in three experiments. Data represent the averages of all experiments (*n* = 3 independent cultures, means ± s.e.m.).

Figure S5. **Effects of the Doa10 deficiency on ER localization of Atg32 truncated variants. (A)** Representative structured illumination microscopy images of *doa10*Δ cells chromosomally expressing Sec63-mRuby2 and one of the truncated Atg32-3×HA-3×GFP variants grown in dextrose medium (SDCA) to $OD_{600}$ = 1.6. Yellow arrowheads indicate Atg32-3×HA-3×GFP overlapping Sec63-mRuby2. Scale bar, 5 μm. DIC, differential interference contrast.

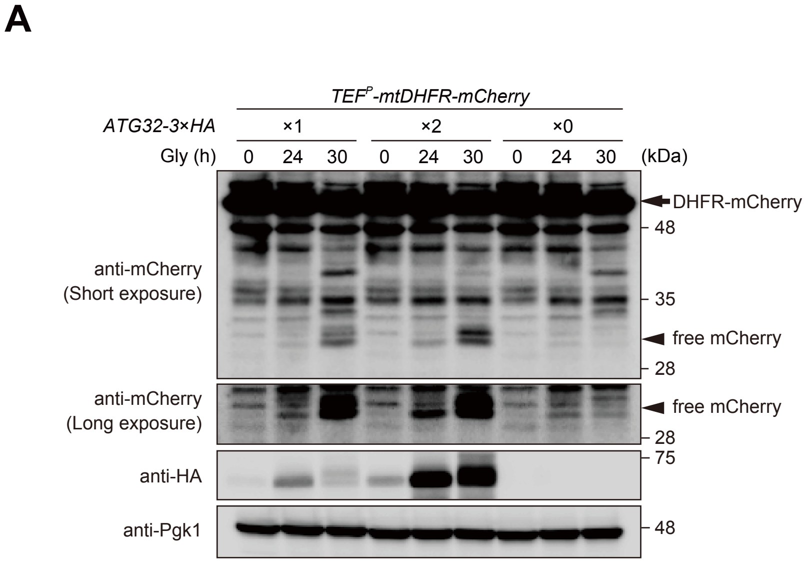
A
TEF^P-mtDHFR-mCherry
ATG32-3×HA
×1
×2
×0
Gly (h)
0 24 30 0 24 30 0 24 30
(kDa)
DHFR-mCherry
48
anti-mCherry
(Short exposure)
35
free mCherry
28
anti-mCherry
(Long exposure)
free mCherry
28
75
anti-HA
anti-Pgk1
48

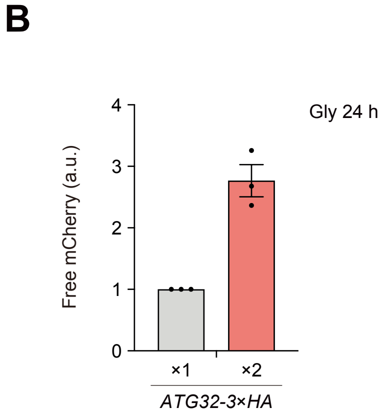
B
Gly 24 h
Free mCherry (a.u.)
0
1
2
3
4
×1
×2
ATG32-3×HA

Eiyama and Onishi et al
Suppl. Fig. 2

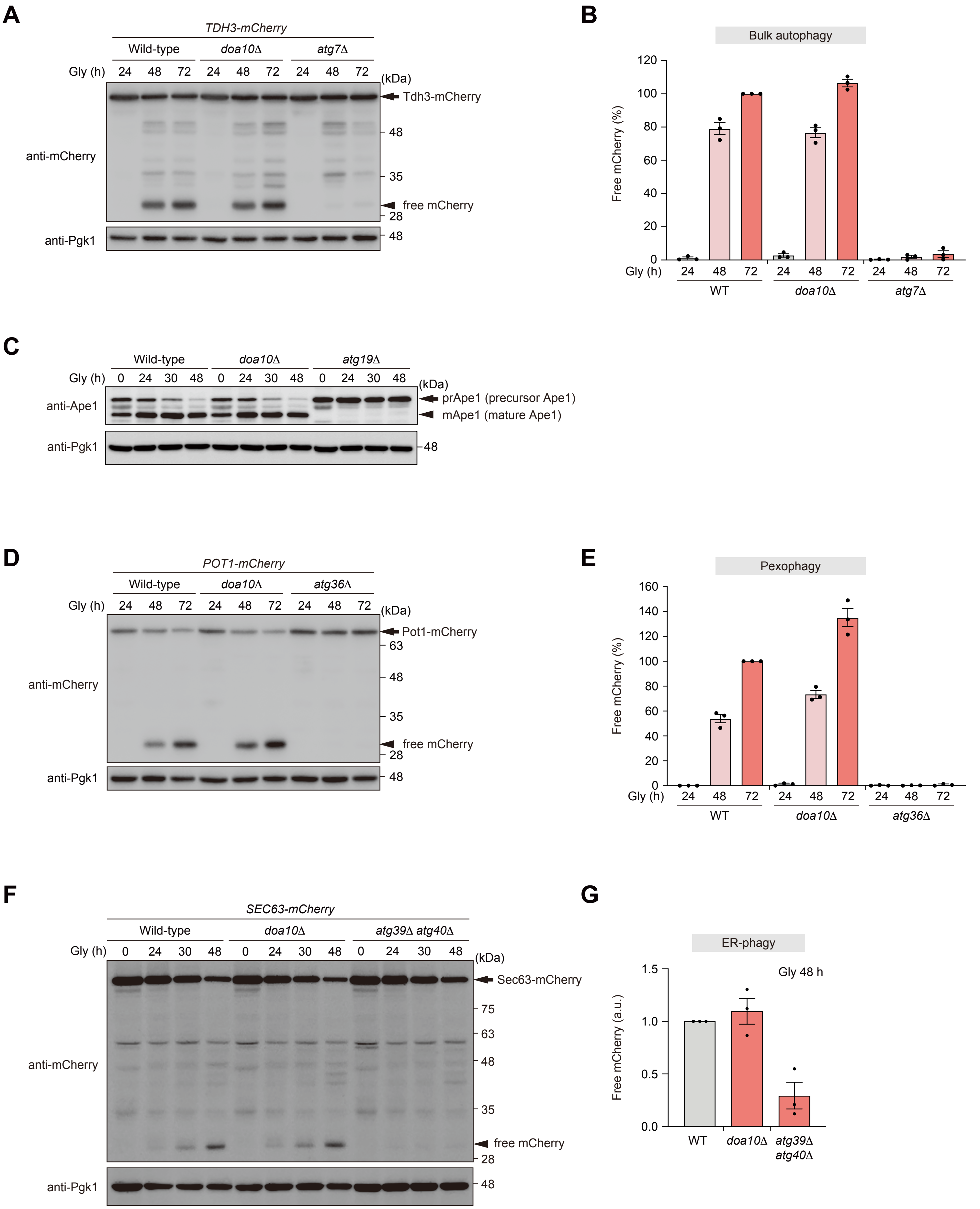

**A**

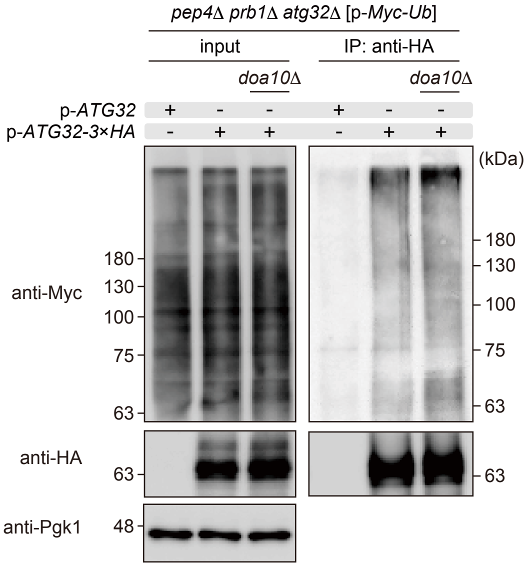
pep4Δ prb1Δ atg32Δ [p-Myc-Ub]
input
IP: anti-HA
doa10Δ
doa10Δ
p-ATG32
p-ATG32-3×HA
+ - - + - -
- + + - + +
(kDa)
anti-Myc
180
130
100
75
63
anti-HA
63
anti-Pgk1
48

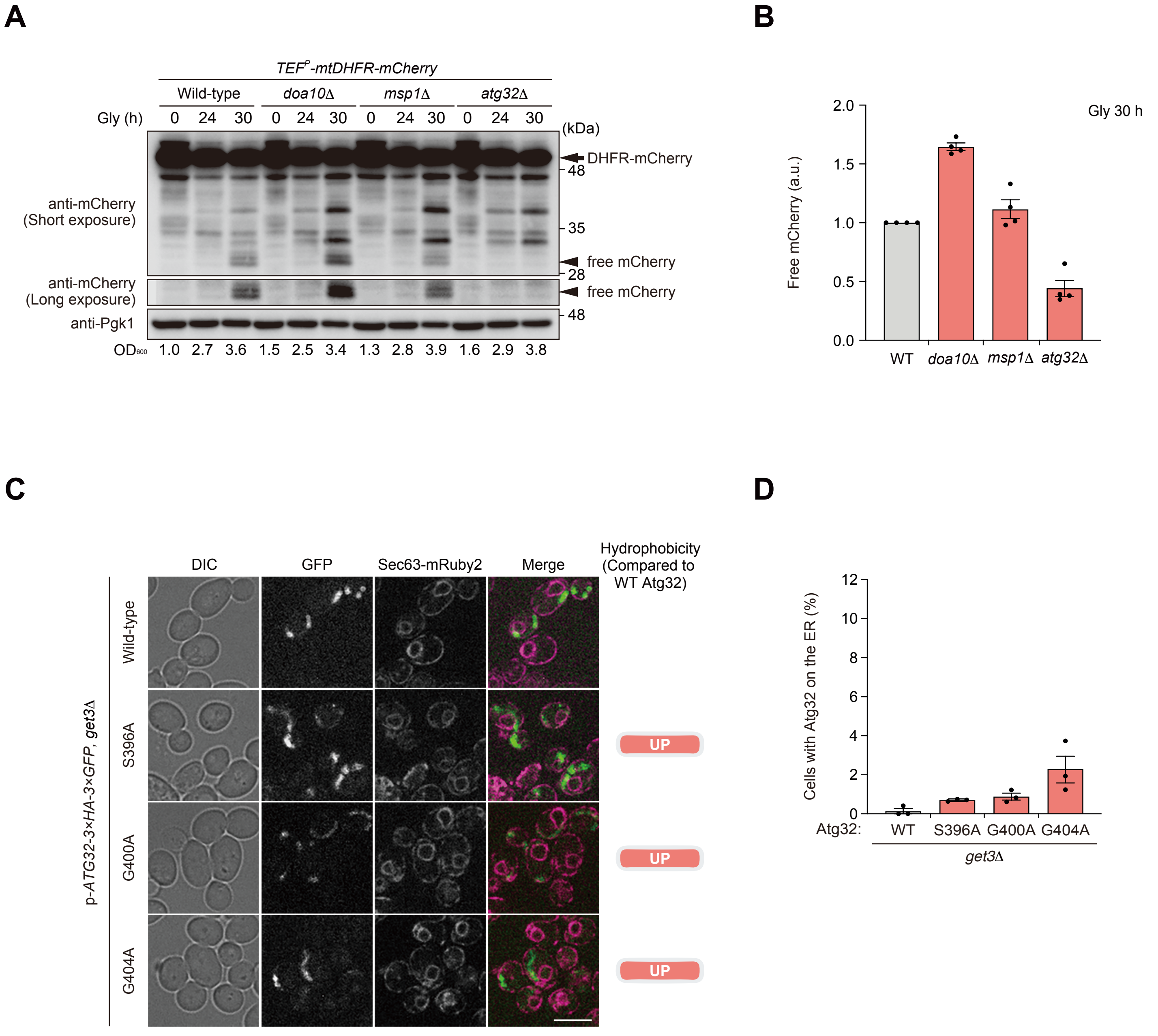
A
TEF^P-mtDHFR-mCherry
Wild-type
doa10Δ
msp1Δ
atg32Δ
Gly (h)
0 24 30 0 24 30 0 24 30 0 24 30
(kDa)
DHFR-mCherry
48
anti-mCherry
(Short exposure)
35
free mCherry
28
anti-mCherry
(Long exposure)
free mCherry
anti-Pgk1
48
OD600 1.0 2.7 3.6 1.5 2.5 3.4 1.3 2.8 3.9 1.6 2.9 3.8
B
Free mCherry (a.u.)
Gly 30 h
2.0
1.5
1.0
0.5
0.0
WT
doa10Δ
msp1Δ
atg32Δ
C
DIC
GFP
Sec63-mRuby2
Merge
Hydrophobicity
(Compared to
WT Atg32)
p-ATG32-3×HA-3×GFP, get3Δ
Wild-type
S396A
G400A
G404A
UP
UP
UP
D
Cells with Atg32 on the ER (%)
12
10
8
6
4
2
0
Atg32:
WT
S396A
G400A
G404A
get3Δ

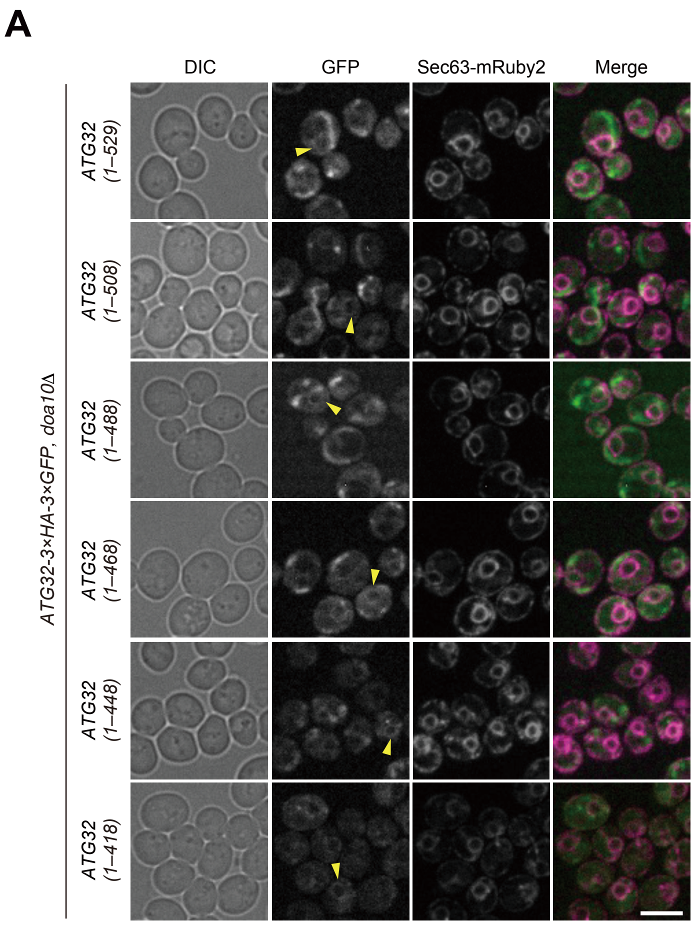
A
DIC
GFP
Sec63-mRuby2
Merge
ATG32-3×HA-3×GFP, doa10Δ
ATG32 (1–529)
ATG32 (1–508)
ATG32 (1–488)
ATG32 (1–468)
ATG32 (1–448)
ATG32 (1–418)
